# Using micro-CT techniques to explore the role of sex and hair in the functional morphology of bumblebee (*Bombus terrestris*) ocelli

**DOI:** 10.1101/433979

**Authors:** David Wilby, Tobio Aarts, Pierre Tichit, Andrew Bodey, Christoph Rau, Gavin Taylor, Emily Baird

## Abstract

Many insects have triplets of camera type eyes, called ocelli, whose function remains unclear for most species. Here, we investigate the ocelli of the bumblebee, *Bombus terrestris*, using reconstructed 3D data from X-ray micro computed-tomography scans combined with computational ray-tracing simulations. This method enables us, not only to predict the visual fields of the ocelli, but to explore for the first time the effect that hair has on them as well as the difference between worker female and male ocelli.

We find that bumblebee ocellar fields of view are directed forward and dorsally, incorporating the horizon as well as the sky. There is substantial binocular overlap between the median and lateral ocelli, but no overlap between the two lateral ocelli. Hairs in both workers and males occlude the ocellar field of view, mostly laterally in the worker median ocellus and dorsally in the lateral ocelli. There is little to no sexual dimorphism in the ocellar visual field, suggesting that in *B. terrestris* they confer no advantage to mating strategies.

We compare our results with published observations for the visual fields of compound eyes in the same species as well as with the ocellar vision of other bee and insect species.

## 1. Introduction

Many insects have two parallel visual systems: a pair of compound eyes that provide spatially-resolved imaging and simple camera-type eyes, known as ocelli. The function of the ocelli – which may vary in number from one to three depending on the species – is unclear for most insects, with implied functionality ranging from simple photostimulatory organs to flight stabilisation and celestial compass-based navigation (e.g. Wellington, 1974; Stange *et al.*, 2002; reviewed in Mizunami, 1994). Ocellar morphology varies tremendously both within species (worker and male honeybees, for example; Ribi *et al.* 2011) and between even closely-related species (Ribi and Zeil, 2018). These differences are thus likely to be driven by selection pressure caused by variation in environment, behaviour and mating strategies as well as potentially reflecting differences in function.

It was assumed for a long time that ocelli were incapable of receiving focused images of the world, as the focal plane of their lenses was generally thought to lie completely behind their retinae (Mizunami, 1994). However, focused regions of visual space have been discovered in various species of dragonfly (Stange *et al.*, 2002; Berry *et al.*, 2007); diurnal and nocturnal species of paper wasp (Warrant *et al.* 2006); the honeybee *Apis mellifera* (Ribi *et al.*, 2011) and the orchid bee *Euglossa imperialis* (Taylor *et al.* 2016). Most recently, detailed investigations of functional morphology of *E. imperialis* ocelli using X-ray micro computed-tomography (micro-CT), which provides a way to investigate 3D visual structures without the slicing damage and potential distortion that accompanies classical microscopy techniques (Baird and Taylor, 2017), has provided the most detailed account of ocellar morphology and fields of view yet (Taylor *et al.*, 2016). The results not only argued against the notion that the ocelli only received under-focused light, but also implied intricate, trinocular navigational functionality that went beyond what had previously been hypothesised.

The buff-tailed bumblebee *Bombus terrestris* (L.) is a widely distributed species of *Bombus*. Its natural habitat spans the temperate regions of Eurasia and it is even considered an invasive species on other continents (Widmer *et al.*, 1998). Like other members of the *Bombus* genus, *B. terrestris* is eusocial, with a caste system consisting of queens, female workers, and male drones. Queens and workers both forage to feed the brood of the queen, while drones only forage for themselves while patrolling for fertile queens. The visual capabilities of *B. terrestris*, have been studied extensively, but these studies have mostly been limited to their compound eyes or visual flight behaviour (e.g. Streinzer & Spaethe, 2014; Foster et al., 2014; Chakravarthi et al., 2016). The few studies that have been conducted on bumblebee ocelli have shown that they have chromatic sensitivity peaking in the ultraviolet range (Meyer-Rochow, 1980) and have suggested that they may play a role in polarization-based navigation (Wellington, 1974), an idea that is supported by the morphology of the ocellar rhabdoms (Zeil et al., 2014). Generally, *B. terrestris* ocelli have been thought to be morphologically similar to the ocelli of *Apis mellifera* (Meyer-Rochow, 1981), although their near-linear external alignment is unusual among insects, most of which sport a more equilateral triangular arrangement (Mizunami, 1994; Streinzer & Spaethe, 2014; Ribi and Zeil, 2018).

With the recent discoveries in hymenopteran ocellar morphology bringing several interesting properties to light, it has become clear that there is still much to be learned about hymenopteran/apid ocellar systems. Studying the species-rich and ecologically varied *Bombus* genus may grant us new insights in hymenopteran ocellar functioning and its relation to hymenopteran visual ecology.

As honeybees and bumblebees face similar sources of ecological pressure, and are closely related (Engel, 1999), it may be expected that the functional morphology of the two species ocelli share many characteristics such as the bipartite ocellar lens and retina found in honeybees (Ribi et al., 2011). The ventral portion of the partitioned ocellar retinae of honeybees, which receives light from the sky, has adaptations that suggest it functions as a skylight detector and/or a celestial compass. The dorsal portion, on the other hand, has adaptations which make it useful as a horizon detector, including a focused view of the world (for the median ocellus). It could also be expected that drone ocelli will invest less in a dorsal view, since drones are less reliant on navigation than workers (Paxton, 2005).

Here, we investigate the functional morphology of the *B. terrestris* ocelli of both workers and males and explore how this relates to their visual ecological needs. To do this, we determined the anatomical, morphological and optical characteristics of bumblebee ocelli using non-destructive micro-CT of preserved samples and the “hanging drop” technique. We then examined how these characteristics shape the fields of view of the different ocelli using computational ray-tracing techniques, for the first time quantitatively investigating the occluding effect of surrounding hairs and comparing the properties of ocellar vision in female worker and male bumblebees. These findings are compared with published data on the fields of view of compound eyes as well as the ocellar fields of view of other flying insects.

## 2. Materials and Methods

### 2.1 Study animals

*Bombus terrestris* workers (female) and drones (male) described in this study were obtained from commercial hives (Natupol, Koppert Biological Systems). This work was carried out in accordance with the Code of Ethics of the World Medical Association (Declaration of Helsinki).

### 2.2 Sample preparation

For this study, six *B. terrestris* individuals (three workers, three drones) were cold anaesthetised, dissected, and prepared for X-ray micro computed-tomography (micro-CT). The lower portion of the head was removed before fixing the sample in a mixture of 3% paraformaldehyde, 2% glutaraldehyde, and 2% glucose in phosphate buffer (pH ~7.3, 0.2 M) for 2 h. Afterwards, samples were washed in buffer and four of these (two workers, two drones) were then fixed in 2% OsO_4_ for 1 h and then dehydrated in a graded alcohol series before being transferred to acetone, and finally embedded in epoxy resin (Agar 100). The external resin was peeled away before scanning. The remaining worker and male heads (with hair intact) were critical-point dried and mounted on pins before scanning.

### 2.3 X-ray micro computed-tomography

Synchrotron micro-CT was conducted at beamline I13-2 of Diamond Light Source in Oxfordshire, UK (Pešić et al. 2013; Rau *et al.*, 2011). Samples were scanned using a polychromatic ‘pink’ beam (5 to 35 keV) of parallel geometry with a 5 mm undulator gap.

The ocelli of 4 embedded samples were imaged in detail. For the workers, 4001 projection images of 0.05 s exposure time were collected by a scintillator-coupled pco.edge 5.5 (PCO AG) detector at equally spaced angles over 180° and at a propagation distance of 120 mm between the sample and the scintillator. For the drones, 8001 projection images of 0.06 s exposure time were collected at a propagation distance of 80 mm. The ocellar scans were conducted with a Castlemaine 4× objective providing 8× total magnification, resulting in an effective isotropic voxel size of 0.81 μm.

The whole dried heads of a worker and a drone were imaged with 2001 projection images of 0.15 s collected at a propagation distance of 80 mm (worker) or 50mm (drone). These scans were conducted with a 1.25× objective providing 2.5× total magnification (2.6 μm voxel size). The radiographic projections acquired from the synchrotron scans were reconstructed, using a filtered back projection algorithm, into 3D volumes using DAWN v1.7 (Basham et al. 2015; Titarenko, 2016).

### 2.4 Analysis of 3D volumes

3D volumes and their pixel intensity range were cropped using Drishti (ANU Vizlab) before they were imported into Amira (FEI). In Amira, volumes were re-sampled to a voxel size of 2 μm (ocellar volumes) or 5 μm (dried head) using a Mitchel filter.

In the ocellar volumes, the lenses, irides, and retinae of the median and left lateral ocelli were labelled manually in every 5th to 10th 2D slice through the volume. Dimensions of the ocelli lenses, irides and retinae were measured in Amira. The retinal thickness was measured on-axis. It was not possible to differentiate between dorsal and ventral retinal regions, as per Ribi *et al.* (2011) and Ribi & Zeil (2018), as individual rhabdoms could not be resolved in our scans. Pigment granules, which are also used to demarcate the dorsal and ventral retinae, did not provide sufficient contrast in X-ray imaging to be visible. The labels for the rest of the volume were acquired through interpolation between the manually labelled slices, manually checking the resulting labels, and smoothing of the resulting volumes. In the dried heads, the external surface was labelled using thresholding functions and the front parts of the ocellar lenses were manually removed (to prevent interference during the ray-tracing analysis). Using different thresholding functions, the hair was separately labelled. Antennae were removed from the labelled volume as it was not possible to account for their position in relation to the eyes (which may vary) and because they are unlikely to have a strong influence on the ocellar fields of view as they are usually pointed frontally during flight. Once complete, the labelled volumes of the ocelli were aligned with respect to the labelled volume of the dried head of the respective caste. This was done through manual rotation, translation and scaling of the labelled ocelli in Amira. The complete head was oriented within the “world” reference frame such that the rear boundary of the compound eyes was vertical and the line between the two lateral ocelli was horizontal (figs. 2C & S1).

### 2.5 Determination of lens optical properties

The ‘hanging drop’ technique, originally described by Homann (1924), was used to determine the optical properties of ocellar lenses. Samples were prepared by collecting ten *B. terrestris* workers that were anaesthetised with CO_2_ and decapitated. After removal of the lower part, the head was submerged in a 70% alcohol solution and kept in a fridge (4° C) for several hours in order to better preserve the ocellar lenses. Afterwards, single ocellar lenses were isolated from the heads and cleaned, with a small piece of cuticle left attached to perform the ‘hanging drop’ technique.

The isolated ocellar lenses were placed on a drop of water on a coverslip, with the interior side in the water, and this coverslip was placed on a rubber O-ring attached with wax to a glass slide, with the drop facing downward (creating a “hanging drop”). The ring was covered in Vaseline to prevent the coverslip from moving. The glass slide was placed under a microscope (Carl Zeiss Axio Scope.A1) with its condenser lens removed. A striped grating printed on transparency film was placed on the microscope lamp. An image of this grating was formed in the water drop by the ocellus, which was viewed through a 40× objective. The back-focal distance (BFD) was determined by measuring the distance necessary to shift the focus of the microscope from the interior surface of the lens to the plane of best focus for the grating. This distance was measured with the micrometre gauge attached to the microscope. To account for observer error, the BFD was measured ten times for every ocellus and averaged. This distance was then corrected to the absolute BFD through multiplication by the refractive index of water (1.33). Of each lens, the focal length was also determined by photographing the focused image with a digital camera attached to the microscope. From this photograph, the spatial wavelength of the image was measured and compared with the spatial wavelength of the actual object.

The focal length, *f*, was determined as

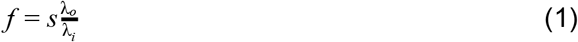

where,

*s*: distance to between the lens and the imaged object (117 mm);

*λ*_*O*_: spatial wavelength of the image of the object; and

*λ*_*i*_: spatial wavelength of the imaged object (7.5 mm).

Refractive indices of the lenses were determined in two different ways. First, the refractive index was estimated by calculating focal length by using the thick lens formula (Born & Wolf, 1999):

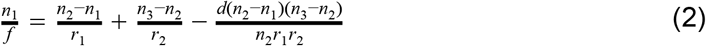

where,

*n*_1_: refractive index of air = 1;

*n*_2_: refractive index of the lens;

*n*_3_: refractive index of the vitreous body = 1.34 (Berry *et al.*, 2007);

*d*: distance separating the inner and outer lens surfaces*

*r*_1_: radius of the outer surface of the lens*

*r*_2_: radius of the inner surface of the lens*

*From the CT volumes used for ray-tracing.

The refractive index of each lens was also determined by systematically varying the refractive index to match the ray-tracer (described in detail below) to the experimental results. Rays were traced through isolated reconstructed lens volumes with different refractive indices in increments of 0.05 between 1.4 and 1.7. The point at which rays converged behind the lens was then determined and compared to measured BFDs. The refractive index which resulted in a BFD closest to the measured BFDs was kept as the refractive index used for further calculations.

Statistical comparisons between the ocelli were made using independent sample Student t-tests or Welch’s t-tests after determining difference in variances using an F-test for equal variances.

### 2.6 Ray-tracing and determination of the field of view

Based on the reconstructed and registered volumes, ray-tracing calculations were performed to determine the field of view of the ocelli. Reconstructed volumes were loaded from image stacks into MATLAB (Mathworks). Voxel-based ray-tracing was then performed using custom-written MATLAB code.

A sphere was created with an infinite radius centred on the centre of the head. The head centre was determined manually during registration, with the centre of the coronal plane defined as the middle between the two antennal bases and the centre on the anteroposterior axis, defined as being directly underneath the median ocellus centre (fig. S1). On this sphere, 2562 point sources were evenly distributed to simulate the panoramic field of view. From each of these point sources, rays were traced across a 7.5 μm grid (separately for each ocellus) such that each ray intersected the front of the lens. The front of the lens was determined by performing a principal component analysis on the 3D lens volume and manually appointing one of the principal components as the principal lens axis, as well as manually determining the polarity along this axis. By tracing rays along the principal axis, the front of the lens was determined as the part where rays hit the lens at the external end of the principal axis. In some cases, this axis had to be manually adjusted such that the lens front was correctly identified. Once a ray intersected with the outer and inner borders of the lens, a quadratic polynomial surface was fitted to the local surface volume, which was used to determine the normal vector to the surface which, in turn, was necessary to calculate the refraction angle of the ray using Snell’s law. The refractive index of the lens was determined as described above. The refractive index of the air was set to 1 and that of the vitreous body to 1.34 (Berry *et al.*, 2007).

A point source was considered to be within the field of view if at least one of the rays from that source entered the retina through the iris aperture, without hitting the iris itself and without being blocked by external features of the head, including cuticle and hair. The point of intersection between the ray and the front surface of the retina was taken as the end point of the ray. The iris aperture was determined using an approach similar to that used for determining the front of the lens, except that the rays traced along the principal axis that did not hit any part of the iris volume were used to create the surface of the iris aperture. The field of view of the lateral ocellus that was not reconstructed (and therefore not used in the ray-tracing) was approximated by mirroring the field of view across the mid-sagittal plane of the median ocellus. This plane was determined by manually adjusting a secondary, perpendicular axis around the previously determined primary axis to create the secondary axis necessary for the creation of the sagittal plane.

For every point source, the number of light rays hitting the retina was taken as a measure of the relative optical power originating from that point source.

Ray-tracing methods were also used to measure the back retinal distance (BRD) for each point source within the field of view. First, an average end point for every point source was calculated by averaging the location of the end points of all rays entering the retina from that point source. The final average end point of the point source was mapped onto the point on the retina front surface that was at the shortest Euclidean distance from the calculated average end point, so that every point source within the field of view was then associated with a voxel on the retinal front surface. Then, normal vectors were calculated for every voxel of the rear surface of the lens by again fitting a quadratic polynomial surface to the surface voxels surrounding the voxel in question and determining the normal vector of the resulting polynomial. Lines based on these normal vectors were then traced through the volume to test whether these lines would cross the vitreous body and the retina without hitting the iris. If this was the case, the Euclidean distance was calculated between the voxel on the lens rear surface and the voxel of the rear retinal surface that crossed the normal of that lens rear surface voxel (i.e. the back retinal distance). This distance was then associated with the voxel of the front surface of the retina that was crossed by the normal. With this information, it was possible to associate the point sources with their respective back retinal distance by cross referencing the end points of the point sources with their corresponding back retinal distance.

## 3. Results

### 3.1 Ocellar Anatomy

The three dorsal ocelli of *Bombus terrestris* are arranged almost linearly on the head (fig. 1A-D) – the wide angle of the triangle formed by the median and lateral ocelli is approximately 170° for both workers and males. Compared to the median ocellus, the orientation of the lateral ocelli (as determined by lens shape and orientation of iris and retina) is rotated around its optical axis by approximately 90° away from the median (fig. 1E,F). Ocelli are surrounded by large tufts of hair both dorsally and ventrally, as well as by hair between the median and lateral ocelli (fig. 1A,B). Reconstructions in 3D reveal that the ocellar retina takes a curved, cupped, “letter C” shape when viewed from above and that the iris, while approximately circular around its outer edge, has an internal extension in the ventral portion which follows the edge of the retina (fig.1G).

**Figure 1:**
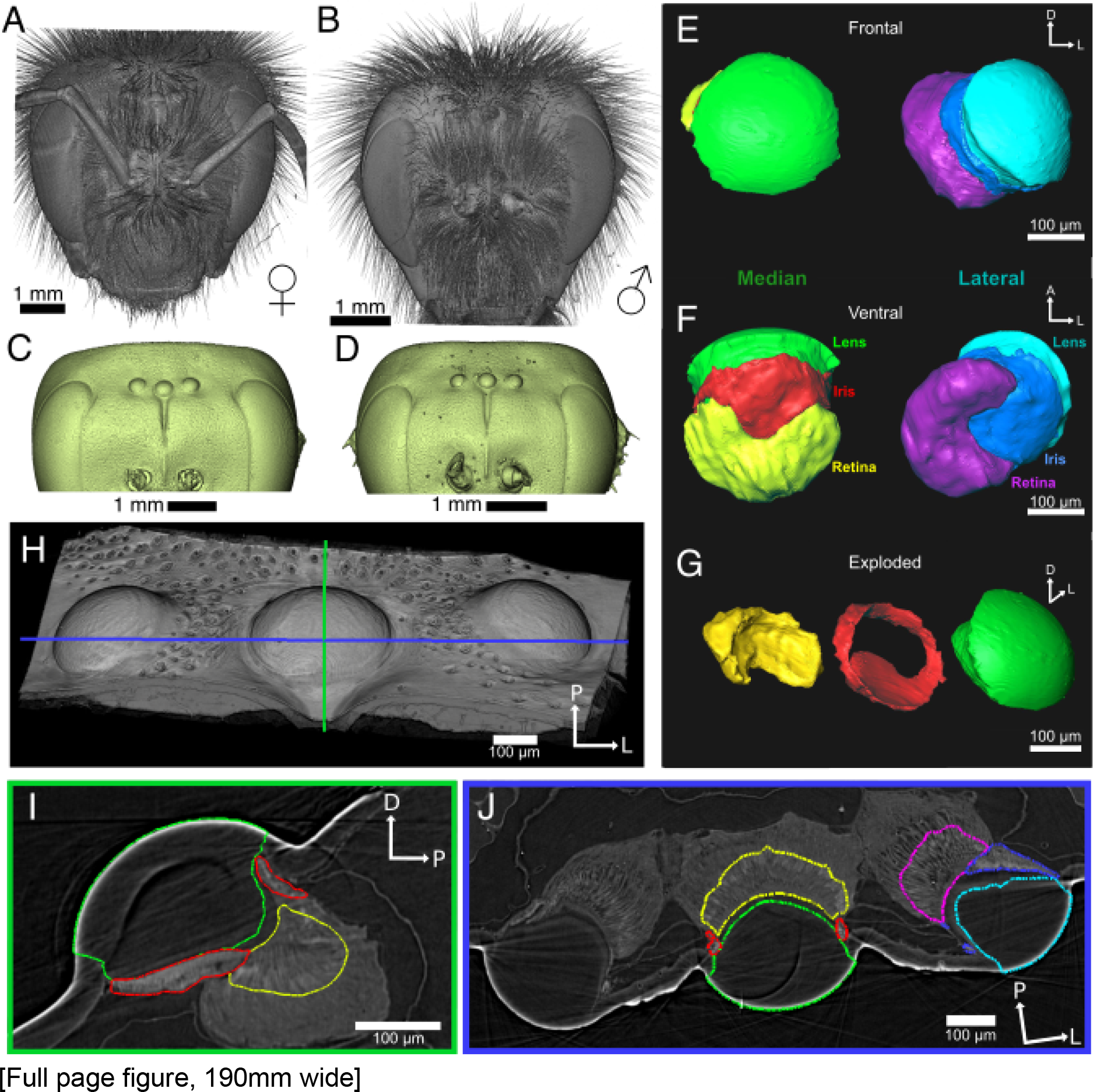
Anatomy of *Bombus terrestris* head and ocelli. A,B: micro-CT reconstructions of worker and male dried heads respectively, showing the dense hair on the head which surrounds the ocelli. C,D: Surfaces of the same heads with the hair and antennae removed computationally, revealing the three ocelli placed dorsally between the compound eyes. E-G 3D volumes of labelled median (left) and left lateral (right) ocelli (E: frontal view, F: ventral view) of a male. The distinct shapes of the iris and retina can be seen, as well as the rotational angle between the median and lateral ocelli. Green & cyan: ocellar lenses, red & blue: irides, yellow and purple: retinae. G: Separate view of the three major parts of the median ocellus, highlighting the curved, elongated, cupped shape of the retina (left), the internal extension of the iris (middle), and the radially asymmetric shape of the back of the lens (right). H-J: Sagittal (I) and transverse (J) sections through the labelled median ocellus of a drone, showing a small vitreous body (between the lens and the retina) as well as the covering of the ventral median portion of the retina by the iris (colour scheme is the same as used above). Section planes are indicated in H. All scale bars: 100 μm. E-J. D: Dorsal, L: Left, A: Anterior, P: Posterior.

The ocellar lenses have an approximately radially-symmetrical, convex frontal surface but the inner surface is asymmetrical and only clearly becomes convex where it meets the small vitreous body (fig. 1G,I,J). Measurements are summarized in table 1. Back retinal distances (BRD) are summarized in table 2.

**Table 1:** Summary of ocellar anatomical measurements. Results are individual measurements from the 4 individuals (2 workers, 2 males) measured.

|  | Worker |  | Male |  |
| --- | --- | --- | --- | --- |
|  | Median | Lateral | Median | Lateral |
| Lens diameter ($\mu\text{m}$) | 264, 292 | 242, 264 | 265, 274 | 274, 247 |
| Lens thickness ($\mu\text{m}$) | 192, 193 | 204, 183 | 177, 207 | 166, 184 |
| Iris aperture width ($\mu\text{m}$) | 226, 255 | 226, 250 | 146, 230 | 220, 226 |
| Retina thickness ($\mu\text{m}$) | 81, 92 | 65, 95 | 88.5, 70 | 90, 79 |

**Table 2:** Summary of back retinal distance measurements. Measurements made from ray-tracing, see text for details of calculations.

| | | BRD [mean ± s.d.] (minimum, maximum) |
| --- | --- | --- |
| Worker | Median | 76 ± 15 $\mu\text{m}$ (min: 43 $\mu\text{m}$; max 134 $\mu\text{m}$) |
| | Lateral | 101 ± 12 $\mu\text{m}$ (min: 21 $\mu\text{m}$; max: 150 $\mu\text{m}$) |
| Male | Median | 84 ± 16 $\mu\text{m}$ (min: 50 $\mu\text{m}$; max: 144 $\mu\text{m}$) |
| | Lateral | 74 ± 14 $\mu\text{m}$ (min: 36 $\mu\text{m}$; max: 117 $\mu\text{m}$) |

### 3.2 Lens Optical Properties

Back focal distances (BFDs) of worker ocellar lenses (as measured with hanging drop) did not differ significantly between the median and lateral ocelli (198 ± 44 μm vs. 184 ± 26 μm respectively, independent samples t-test assuming equal variances: t(6,6) = 0.65, p = 0.53), and neither did the focal length (341 ± 84 μm vs. 334 ± 87 μm, independent samples t-test assuming equal variances: t(8,6) = - 0.16, p = 0.88). Determination of lens refractive index, *n*_*l*_, through match of BFDs in the ray-tracer resulted in a *n*_*l*_ value of 1.5 for all 4 median ocelli and 2 lateral ocelli, and an *n*_*l*_ value of 1.45 for 2 lateral ocelli. Determination of *n*_*l*_ with the thick lens formula (rounded to the nearest .05) resulted in a *n*_*l*_ value of 1.55 for all median ocelli and an average of 1.58 for lateral ocelli. Based on these results, *n*_*l*_ was set to 1.5 in the ray-tracing simulations. It was assumed that the refractive index of male ocelli is similar to those of workers. BFDs were substantially longer than the BRDs, indicating that it is unlikely that images are focused anywhere on the retina.

### 3.3 Fields of View

Ray-tracing calculations were used to model the field of view (FoV) of worker and male bees, accounting for the occlusion of rays by hairs around the ocelli (fig. 2). The median ocellar FoV is centred around the sagittal plane (azimuth = 0°), whereas the lateral FoV falls entirely to the left or right of this plane. All ocelli view the horizon (elevation = 0°). Without accounting for hair occlusion, the lateral ocellar FoV continues laterally and views part of the posterior-dorsal field. There is a binocular overlap between the two lateral ocelli dorsally, though this does not always include the median ocellus for the specimens tested (fig. 2E,G,I,K and see fig. S2).

**Figure 2:**
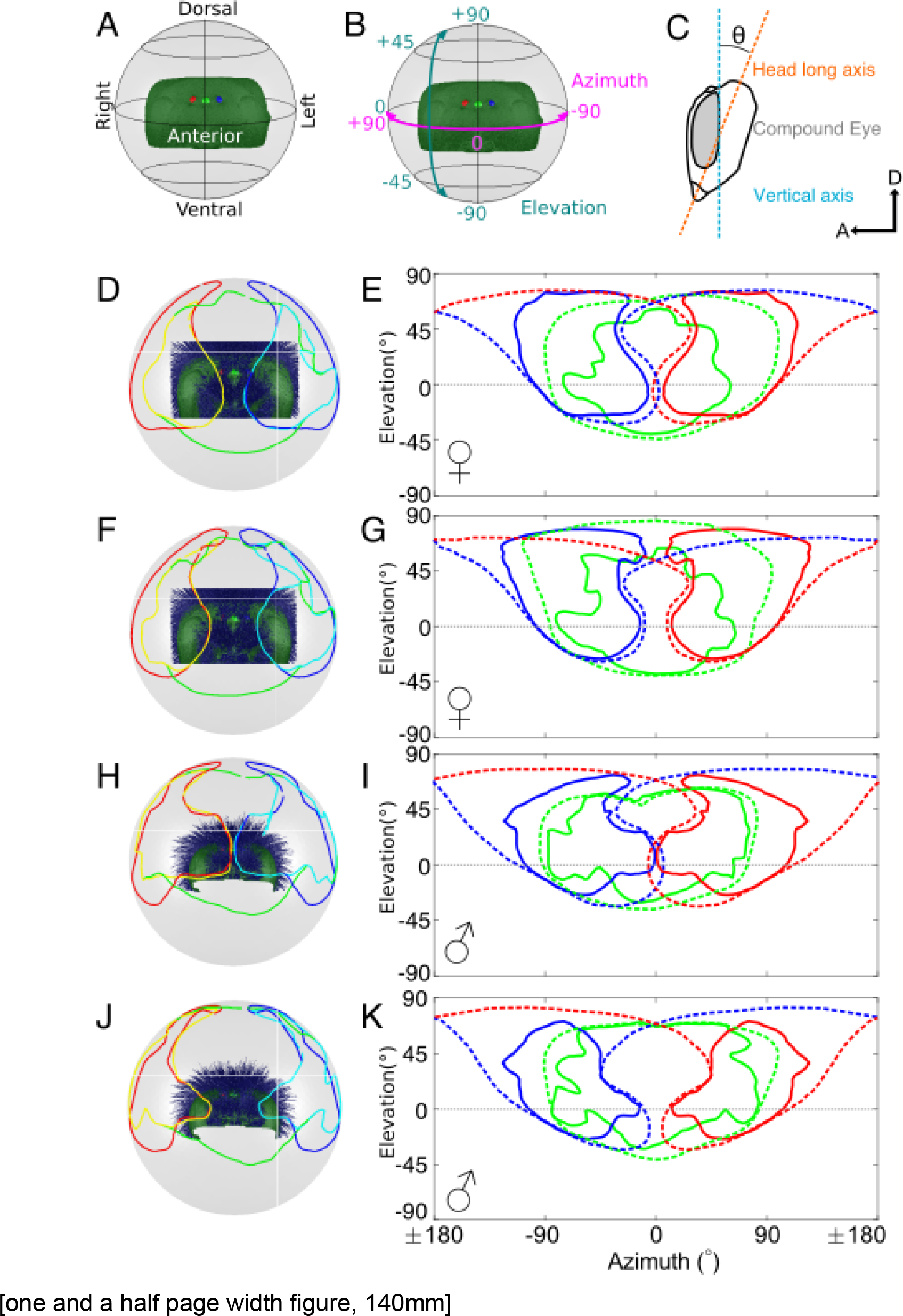
Visual fields of ocelli of worker and male bumblebees. Boundaries of the simulated visual fields are shown in spherical and equirectangular projections. A: Illustrates how the head (dark green) and ocelli (light green: median; red: right lateral; blue: left lateral) are oriented in the spherical projection plots (D,F,H,J). B: Illustrates the relationship between spherical projections and coordinates of the equirectangular projection plots (E,G,I,K). C: Orientation of the head with respect to pitch angle, θ, relative to the long axis of the head. In each of the plots D-K, solid lines enclose a visual field occluded by hair (shown in dark blue) for the three ocelli. Dashed lines in plots E,G,I,K indicate the un-occluded field of view. D,E; F,G: Two different worker individual ocelli registered onto another worker dried head. H,I; J,K: Two different male individual ocelli registered onto another male dried head. Cyan and yellow lines in D,F,H,J represent regions of the visual field viewed by both the lateral and median ocelli.

Little, if any, sexual dimorphism was observed in the ocellar FoV and the overall size and orientation of each visual field was similar. Total visual field was slightly smaller in the males studied (fig. 3).

**Figure 3:**
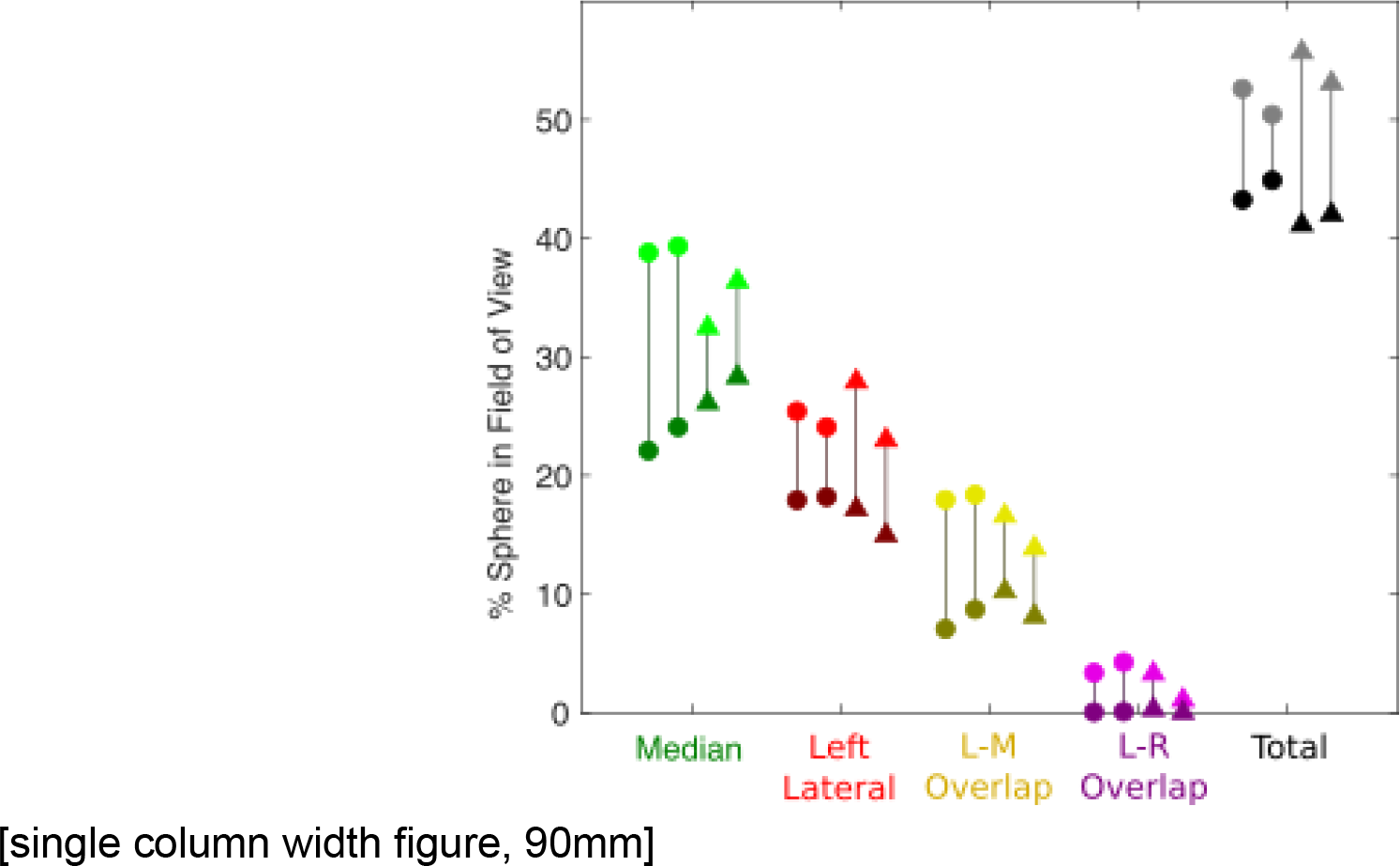
Percentage of “World Sphere” inside Field of View for each ocellus and overlap between ocellar fields of view. Lighter, upper symbols show values for calculations excluding hairs and darker, lower, darker symbols show values for those including hair occlusion, lines connect values for the same individual. Workers are shown as circles and males as triangles. L-M overlap is the overlap in field of view between the left lateral and median ocelli. L-R overlap is the overlap in field of view between the left and right lateral ocelli. Total is the unique field of view of all three ocelli.

In all cases tested, hairs restrict the FoV of the ocelli. Median ocelli were occluded primarily in the lateral and dorsal directions and lateral ocelli mostly in the dorsal FoV. The ocelli whose FoV was most occluded by hair were worker median ocelli and male lateral ocelli (figs. 2 & 3). The total ocellar FoV is predominantly anterior-dorsal, with some lateral ocellar FoV extending above an azimuth of 90°. The lateral ocellar FoV extends further dorsally than the median ocellus.

Binocular overlap between the median and left lateral ocelli was around 10% of the total viewing sphere, though this was reduced from 14-18% by hair occlusion (fig. 3). Before occlusion, there was little overlap between the two lateral ocelli, though this was reduced to nil by the hairs (fig. 3). There was no trinocular overlap calculated between the three ocelli. In all cases, a gap between the two lateral ocellar fields of view was observed in the dorsofrontal region (fig. 2D-K).

As a measure of the relative optical power distribution, the number of rays which struck the retina from each source were counted and this number normalised across all of the rays that reached the retina in all samples. It can be seen (fig. 4 and see fig. S3) that the optical power for the median ocellus is slightly greater for a vertical band in the centre and a horizontal band at the ventral part of the FoV. Variation in the power distribution of the lateral ocellus is less pronounced, but is greatest at the ventral and rearmost parts of its FoV. Relative optical power for the males was typically lower than for the females, but a similar distribution pattern was seen. In all cases, occlusion by hairs reduced the optical power across the whole FoV.

**Figure 4:**
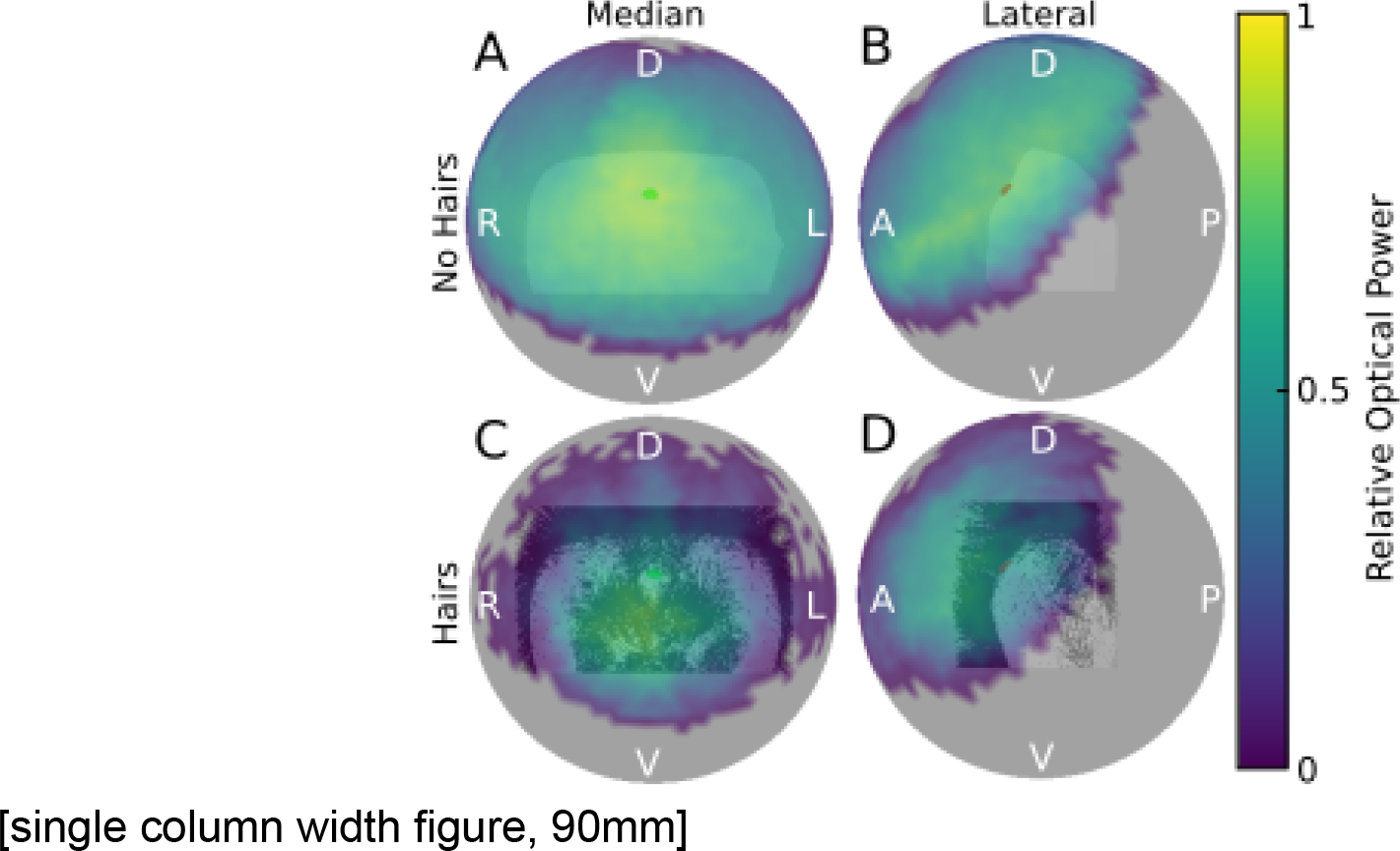
Relative optical power for bumblebee worker ocelli. Relative optical power (normalised number of simulated rays from each point source which strike the retina) of the visual field of the median (A,C; frontal view) and left lateral (B,D; lateral view) ocellus corresponding to the same individual shown in fig. 2D,E. Note that the color map is plotted transparently on a sphere around the head surface (as per the fields of view in Fig. 2). A,B: No occlusion. C,D: Occluded by hairs.

## 4. Discussion

Bumblebee ocelli are unusual in two ways: firstly, their relative positioning to one another is almost linear (fig. 1A-D; e.g. Streinzer & Spaethe, 2014), whereas most hymenopteran ocelli are placed in an arrangement closer to an equilateral triangle (e.g. Ribi & Zeil 2018); secondly, they are surrounded by hairs that restrict their field of view (as shown here), whereas the ocelli of locusts, blowflies, dragonflies etc. have no substantial occlusion (Taylor, 1981; Schuppe & Hengstenberg, 1993; Stange *et al.* 2002). Other than this, *B. terrestris* ocelli retain the major features of the majority of insect ocelli such as their wide fields of view, under-focused optics, dorsal positioning and high optical sensitivity.

### 4.1 Arrangement of the Ocelli

The unusual, near-linear arrangement of *B. terrestris* ocelli (fig. 1A-D) is potentially an adaptation to provide a forward-looking visual field for each ocellus while remaining in a dorsal location on the head. For example, the ocelli of the desert locust (*Schistocerca gregaria*) and dragonflies (*Hemicordulia tau* & *Orthetrum caledonicum*) are also forward-looking, but are positioned in a triangular formation on the front of the head, with fields of view centred on the horizon (Taylor, 1981; Stange *et al.* 2002; Ribi & Zeil 2018; reviewed in Goodman, 1970). In contrast, the ocelli of the blowfly (*Calliphora erythrocephala*) and the orchid bee (*Euglossa imperialis*, fig. 6), which are arranged in a triangle, are dorsally-positioned on the head and have approximately dorsally-centred fields of view (Schuppe & Hengstenberg, 1993; Taylor *et al.* 2016; Ribi & Zeil 2018). It is unclear exactly what benefits arise from these different ocellar orientations but we speculate that this may be related to differences in the visual information available in the habitat and/or reflects differences in their function.

### 4.2 Ocellar Fields of View

The unoccluded FoVs of the ocelli are large and cover over 50% of the world when combined (fig. 3). The median ocellus looks directly forwards and at the horizon, with its FoV extending almost to the dorsal pole (fig. 2E,G,I,K). Lateral ocellar FoVs look sideways and dorsally, having a considerable binocular overlap between them near the dorsal pole. The overall ocellar FoV is anterior rather than dorsal, which would suit the needs of horizon-based head stabilisation during flight in an open habitat.

Similar to the ocellar fields of view of locusts and dragonflies, *B. terrestris* ocelli have visual fields which encompass the horizon (fig. 2D-K). However, in this study, the head pitch angle was set such that the rear boundary of the compound eyes was vertical (fig. 2C) - in line with the analyses in Taylor *et al.* (2016 & preprint). In *B. terrestris* this boundary is tilted around 10-18° further forward than the “long axis” of the head which has been used in behavioural studies to examine head position during flight and landing (Reber *et al.* 2016). To our knowledge, there are no published data for the head pitch angle of bumblebees during free flight (i.e. other than during landing) so this angle may vary between 0° and 40° backward from vertical. As such, data here can be thought of as being correct for the case of around 10-18° backward head pitch. For a head pitch of 0°, the ocellar fields of view would be lowered slightly towards the horizon.

### 4.3 Sexual Dimorphism

We found little evidence for sexual dimorphism in the ocellar anatomy or visual fields of *B. terrestris*. This is somewhat surprising due to the large differences between the behavioural ecology of workers and males. Females (workers) carry out foraging tasks – requiring them to navigate repeatedly between food sources (flowers) and their nests – whereas males (drones) solely seek female queens to mate with. Within *Bombus* species, there are two major mating strategies among males - perching and visually targeting queens; or using scent-marking and patrolling a defined route to attract them. Streinzer & Spaethe (2014) noted that males of *Bombus* species which employed a perching mating strategy had enlarged compound eyes with enlarged facets in the frontal regions compared to workers of the same species. On the other hand, males of scent-marking species, including *Bombus terrestris*, showed little sexual dimorphism in the compound eyes. Our observations suggest that a similar pattern is found in the ocellar apparatus of *B. terrestris*, suggesting that enlarged or specialized ocellar visual fields do not confer a selective advantage in mating. It is also possible that the ocelli play a more general role in flight control such that they are necessary for basic behaviours such as head stabilisation and orientation that all individuals, be they male or female, must perform to remain aloft.

### 4.4 The Effect of Hairs on Ocellar Vision

To our knowledge, this is the first study to quantitatively examine the occluding effect that structures on the head, such as prominent tufts of hair, have on the field of view. Ribi *et al.* (2011) and Hung & Ibbotson (2014) estimate that hair around honeybee ocelli likely obscures the FoV of all three ocelli in the dorsal direction and shields the median ocellus from bright overhead light.

We find that for *B. terrestris*, the FoV of ocelli are indeed limited by hair in the dorsal direction and that the overlap between ocellar FoV is reduced as a result of hair occlusion. However, the hairs do not drastically alter the median ocellar FoV in the dorsal direction for all of the specimens tested and there is still considerable overlap between the FoV of the lateral and median ocelli despite its reduction. The overlap between the two lateral ocelli is completely eliminated by the hairs. Further, our modelling shows that the optical power reaching the ocellar retina is reduced by the hairs (figs. 4 & S3). It is possible that the hairs do play an important role in limiting and regulating the fields of view of the different ocelli. Interestingly, not all Bombus species have the same amount of hair on their heads, with Arctic species having much thicker hair than tropical species, which have very little hair (personal observations). Exploring the effect of hair in different species may therefore help to shed light on the potential role of hair on ocellar function. Our findings also highlight the importance of exploring the effect of any structures on the head (such as horns in beetles) on the visual field when investigating animal vision.

### 4.5 Relationship with Compound Eyes

A feature shared by all of the specimens tested here is a space in the dorso-frontal region (azimuth ≅± 20°, elevation ≅ 20-50°) which is not viewed by the lateral ocelli (fig. 2E,G,I,K; fig. 5). However, when compared with the visual fields of the compound eyes of workers (Taylor *et al.* preprint), it becomes clear that this area is viewed by the median ocellus and both compound eyes. It is not clear what, if any, the function of this overlap between the two visual systems is. Further work investigating whether information from regions in space where both visual systems overlap is combined neurally would help to answer this question. A potential role of the ocelli as having a supporting function for the compound eyes is also suggested by our observation that the FoV of the lateral ocelli extend dorsolaterally past the field of view of the compound eyes (fig. 5). This area could potentially respond to relevant stimuli in this area, this helping to extend the visual reach of the compound eyes.

**Figure 5:**
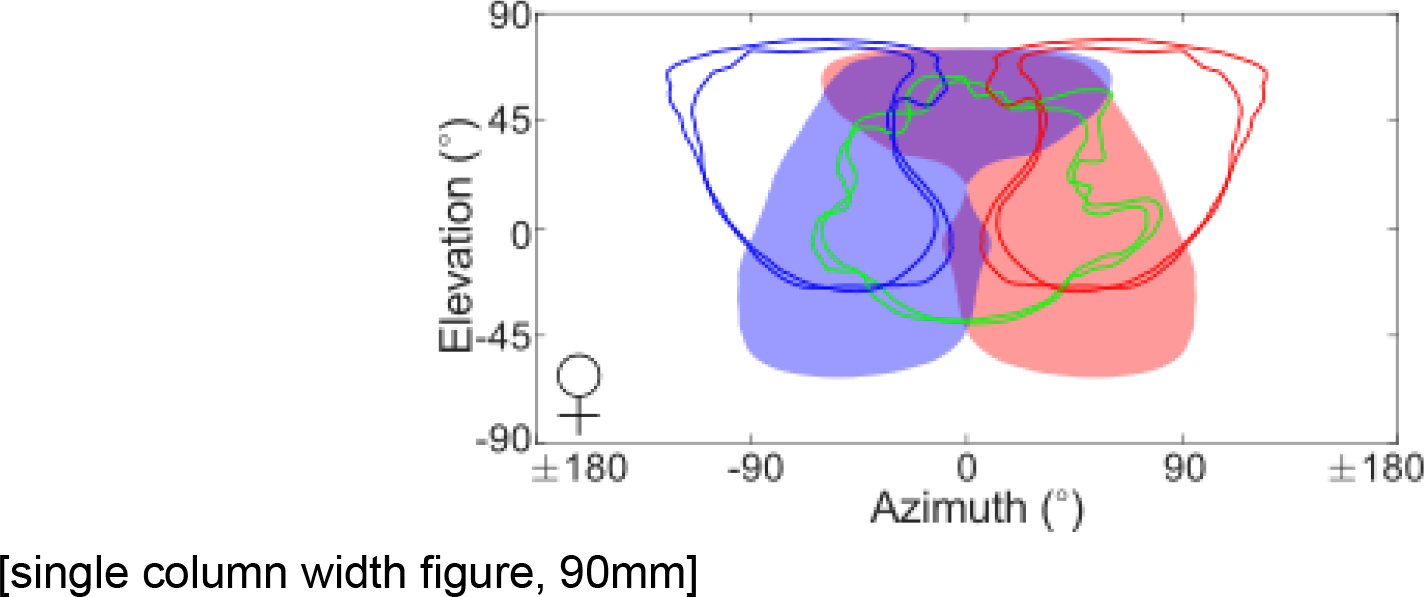
Fields of view of worker bumblebee (hair occluded) ocelli and compound eyes. Lines show the hair occluded fields of view of both *B. terrestris* worker ocelli (as in fig. 2). Shaded areas show the fields of view of typical left (blue) and right (red) compound eyes as calculated by Taylor *et al.* (preprint).

### 4.6 Comparison with other bee species

As social, central-place foraging hymenopterans, honeybee (*Apis mellifera*) and bumblebee (*B. terrestris*) workers are often considered to have similar visual systems. However, our results show that there are numerous striking differences in their ocelli. Worker honeybee ocelli are arranged in a triangle on the top of the head, and on the front of the head in males/drones and, with the exception of the dorsal retina of the median ocellus, receive under-focused images (Ribi *et al.* 2011). Similarly, honeybee ocelli are surrounded by hairs, which likely limit their field of view such that the worker median ocellus looks forward, whereas the lateral ocelli look sideways and upwards (Hung & Ibbotson, 2014). From observations of external morphology, Ribi *et al.* (2011) estimate that drone ocelli all appear to point forwards, with the lateral ocelli partially occluded on the sides by the extremely large compound eyes and on their dorsal side by a tuft of hair. Hung & Ibbotson (2014) suggest that the worker honeybee ocellar dorsal retina samples the horizon, whereas the ventral retina samples the sky. However, no quantitative study has been undertaken on the visual fields of honeybee ocelli and very little is known about how this compares to the male visual system.

On an internal morphological level, parallels with bumblebees are less apparent. Whereas the honeybee ocellar lens sports a cusp in the outer surface, separating dorsal and ventral sides of the ocellar lens front, there is no such partition in the bumblebee ocellar lens. The inner surface of the ocellar lens of the honeybee is radially asymmetrical, but this asymmetry appears to be far greater in the bumblebee. The vitreous body of the honeybee ocellus is more extensive relative to ocellar lens size than that of the bumblebee ocellus. The ocellar retina of the honeybee, while not as stretched as in the bumblebee, is similarly thick as the bumblebee ocellar retina. However, the clear distinction that is made between the dorsal and ventral portions of the retinae cannot reliably be made in the bumblebee due to limitations in our ability to image pigment granules. However, while it has not previously been commented upon, the shape of the iris of honeybee ocelli *does* appear to have a similar internal extension as we found for bumblebee (Ribi *et al*., 2011, Ribi & Zeil 2018).

Another bee species which has previously been studied in terms of its ocellar vision is the orchid bee (*Euglossa imperialis*). Male orchid bees have their three ocelli arranged in a triangle and positioned on the dorsal part of the head, with fields of view that each fill most of the dorsal hemisphere and that overlap to give a region of trinocular vision and have regions of image focus (Taylor *et al.* 2016). Taylor et al. (2016) hypothesised that this is an adaptation to their visual habitat of dense rainforest, meaning that there is often no clear horizon and as such the ocelli look directly upwards (fig. 6) to obtain visual information including analysis of the skylight polarization pattern. As orchid bees have little hair on their heads, the visual fields of their ocelli are not occluded by hair. These properties of orchid bee ocelli are clearly quite distinct from what we see in bumblebees, potentially reflecting differences in habitat, behavioral ecology and/or function.

**Figure 6:**
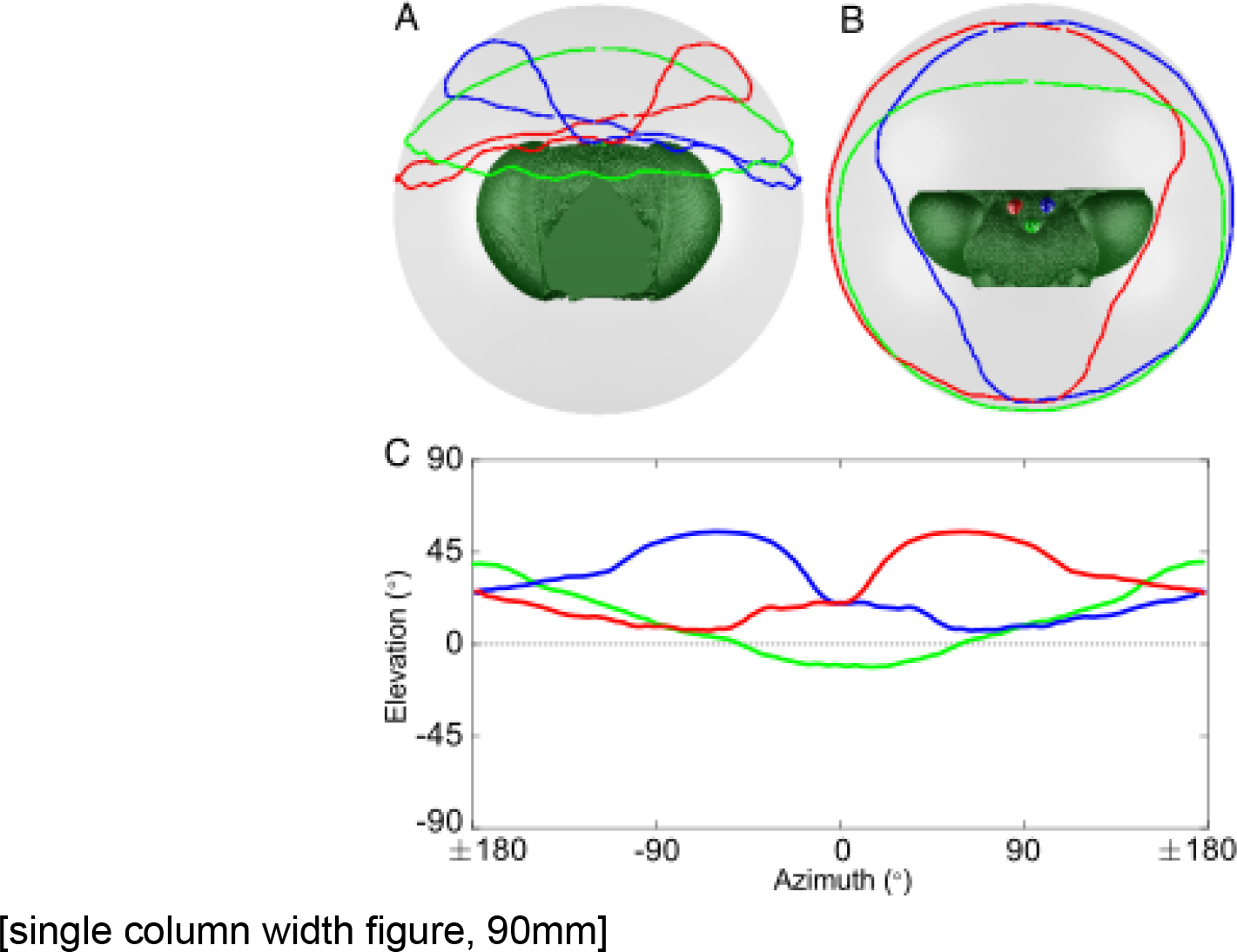
Ocellar fields of view of the male orchid bee, *Euglossa imperialis*. A) Frontal view. B) Dorsal view. C) Equirectangular projection - lines indicate the lower boundary of the field of view. (replotted with permission from Taylor *et al.* 2016). As above, light green: median ocellus; red,blue: right,left lateral ocelli respectively.

### 4.7 Role of the Iris

Many ocelli contain a pigmented iris in a ring-shape between the lens and the retina. Honeybees, orchid bees and bumblebee ocelli also possess an iris (Ribi *et al.*, 2011; Ribi & Zeil 2018; Taylor *et al.*, 2016) and here, we show using 3D reconstruction that in *B. terrestris* the iris, whilst roughly circular around its outer edge, has a curved, “c-shaped” aperture, following the shape of the retina itself (fig. 1G). It may be that in bumblebees the shape of the iris is also variable, though no variations in shape have been noted between the four specimens in this study, and may be involved in light adaptation as in dragonfly ocelli (Stavenga *et al.*, 1979). A similar, curved iris has been found in *Triatoma infestans*, though this does not constrict in response to light (Insausti & Lazzari, 2000).

In the median *B. terestris* ocellus the iris covers the most ventromedial part of the retina, similar to the iris of the dragonfly (Stavenga *et al.*, 1979). This, together with the hairs dorsal to the ocellus, shields the most ventral portion of the retina from direct sunlight. Skylight is, however, important for the proposed function of bumblebee (Wellington, 1974), orchid bee (Taylor et al., 2016) and honeybee ocelli (Ribi et al., 2011; Ribi & Zeil 2018; Ogawa *et al.*, 2017), which likely make use of the celestial polarization pattern, and none of their respective ocellar irides noticeably cover the ventral portion of their retinae. Further investigations into the function of the ocelli and the structure of their irides are necessary for understanding their role in vision in these insects.

## 5. Conclusions

We used X-ray micro computed-tomography and computational ray-tracing to investigate the structure and fields of view of bumblebee (*Bombus terrestris*) ocelli. We make comparisons between workers (female) and drones (male), finding little sexual dimorphism between their ocelli. Our measurements of the ocellar morphology and lens refractive index indicates that, like many insect ocelli, those of *B. terrestris* are under-focused. Using ray-tracing simulations, we calculate that median ocelli look forwards and at the horizon; and that lateral ocelli have a field of view which is directed somewhere between forwards and sideways, but also extends upwards. Further, we have, for the first time, quantitatively tested the occluding effects of hairs on ocellar vision, showing that hairs do shield the ocellar fields of view, reducing binocular overlap and blocking the dorsal field of view of the lateral ocelli.

## Acknowledgements

We would like to thank Carina Rasmussen, Eva Landgren, and Ola Gustafsson for facilitating sample preparation and providing access to the Microscopy Facility at the Department of Biology, Lund University and Viktor Håkansson for assistance preparing samples as well as Qiang Tao, Rajmund Mokso, and Kazimir Wanelik for providing assistance during micro-CT imaging.

## Supplemental Information

**Figure S1:**
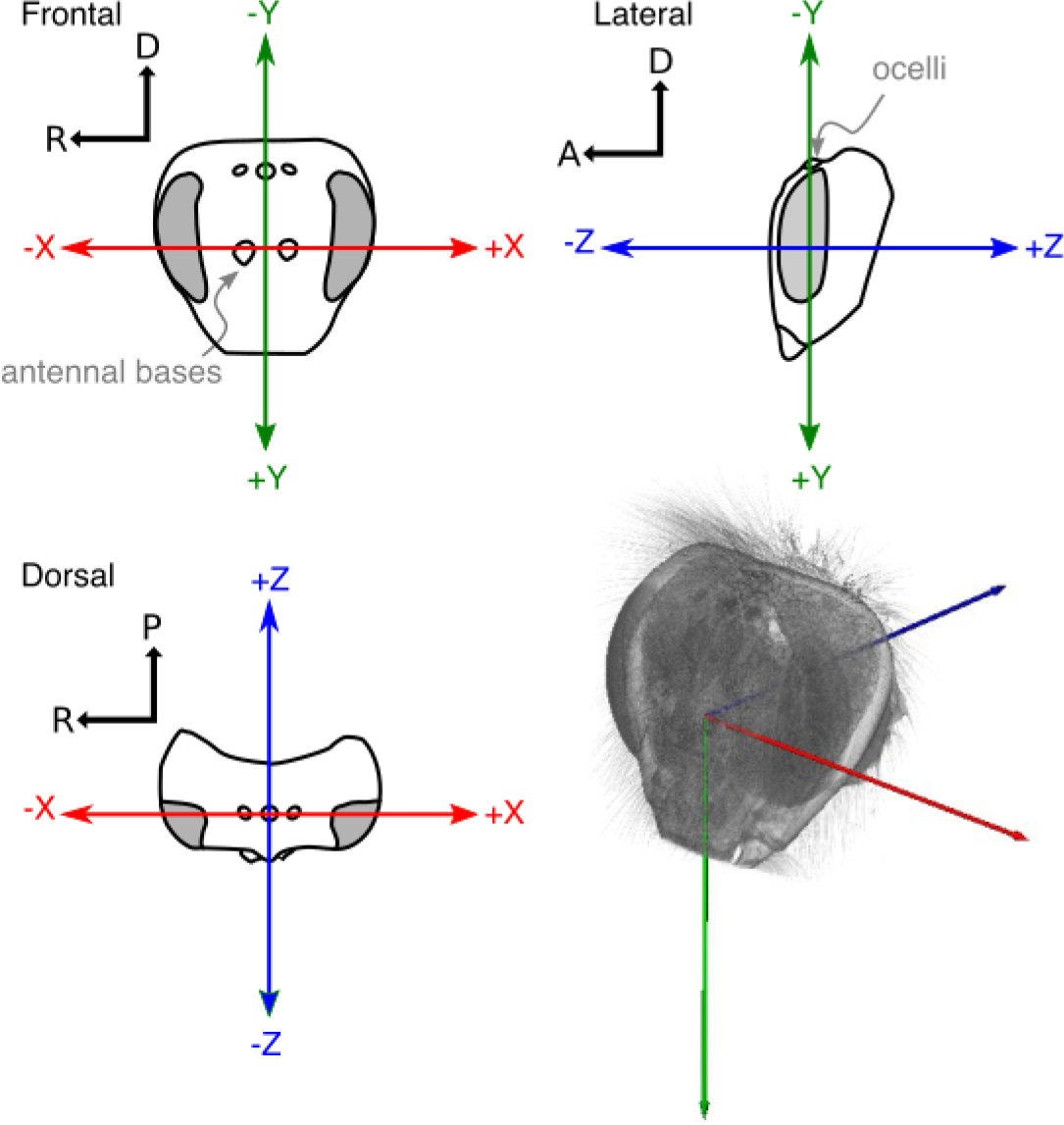
Orientation and alignment of the head for ray-tracing simulations. The head was aligned such that the tops of the compound eyes and lateral ocelli were aligned with the horizontal (X) axis; the Z-axis directly ran through the centre between the two antennal bases; and the vertical (Y) axis ran through the centre of the median ocellus. The head pitch-angle was set such that the rear edge of the two compound eyes was parallel to the vertical axis (also see fig. 2C in the main text). D: dorsal; R: right; A: anterior; P: posterior.

**Figure S2:**
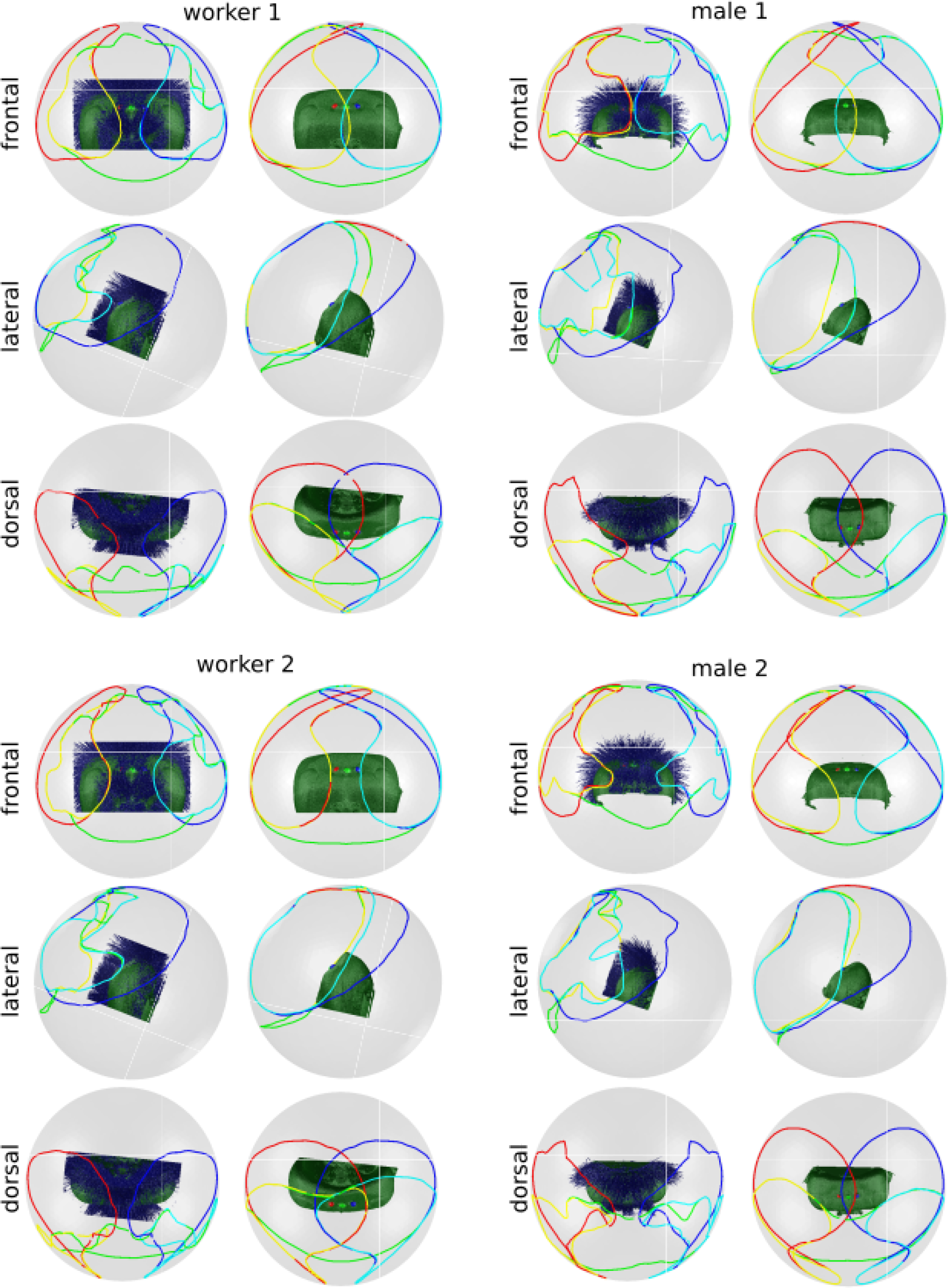
Spherical projections of ocellar fields of view. As per figure 2 in the main text. Worker 1 corresponds to fig. 2D,E; worker 2 - fig.2 F,G; male 1 - fig. 2 H,I; male 2 - fig. 2 J,K.

**Figure S3:**
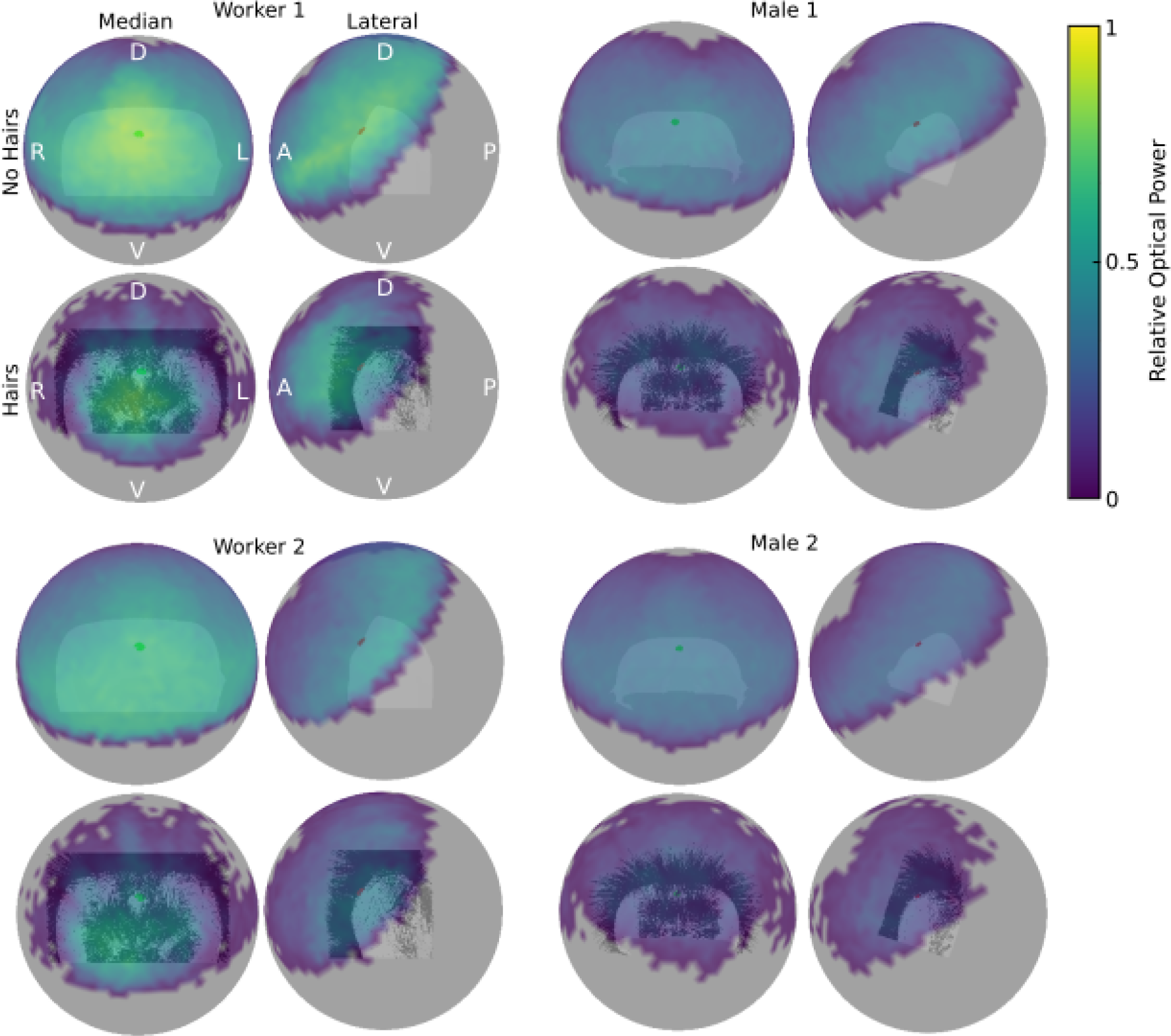
Relative optical power for all ocelli tested. As per figure 4 in the main text. Worker 1 corresponds to fig. 2D,E; worker 2 - fig.2 F,G; male 1 - fig. 2 H,I; male 2 - fig. 2 J,K.

